# *Trpc6* gain-of-function disease mutation enhances phosphatidylserine exposure in murine platelets

**DOI:** 10.1101/2022.02.01.478727

**Authors:** Kimber L. Boekell, Brittney J. Brown, Brianna E. Talbot, Johannes S. Schlondorff

## Abstract

Platelets enhance coagulation by exposing phosphatidylserine (PS) on their cell surface in response to strong agonist activation. Transient receptor potential channels, including TRPC6, have been implicated in the calcium influx central to this process. Here, we characterize the effect of a *Trpc6* gain-of-function (GOF) disease-associated, and a dominant negative (DN), mutation on murine platelet activation. Platelets from mice harboring *Trpc6^E896K/E896K^* (GOF) and *Trpc6^DN/DN^* mutations were subject to *in vitro* analysis. *Trpc6^E896K/E896K^* and *Trpc6^DN/DN^* mutant platelets show enhanced and absent calcium influx, respectively, upon addition of the TRPC3/6 agonist GSK1702934A (GSK). GSK was sufficient to induce integrin αIIbβ3 activation, P-selection and PS exposure, talin cleavage, and MLC2 phosphorylation in *Trpc6^E896K/E896K^*, but not in wild-type, platelets. Thrombin-induced calcium influx and PS exposure were enhanced, and clot retraction delayed, by GOF TRPC6, while no differences were noted between wild-type and *Trpc6^DN/DN^* platelets. In contrast, Erk activation upon GSK treatment was absent in *Trpc6^DN/DN^*, and enhanced in *Trpc6^E896K/E896K^*, platelets, compared to wild-type. The positive allosteric modulator, TRPC6-PAM-C20, and fluoxetine maintained their ability to enhance and inhibit, respectively, GSK-mediated calcium influx in *Trpc6^E896K/E896K^* platelets. The data demonstrate that gain-of-function mutant TRPC6 channel can enhance platelet activation, including PS exposure, while confirming that TRPC6 is not necessary for this process. Furthermore, they suggest that *Trpc6* GOF disease mutants do not simply increase wild-type TRPC6 responses, but can affect pathways not usually modulated by TRPC6 channel activity, displaying a true gain-of-function phenotype.

## Introduction

Influx of calcium ions into the cytoplasm is a versatile signaling event utilized by most cell types, including platelets.[1] Through variation in the location, duration and magnitude of calcium influx, and integration with other signals, calcium signaling can influence a wide range of cellular responses. In platelets, calcium signaling is involved in multiple aspects of activation, including granule release, phosphatidylserine exposure, and shape change.[2] Several different calcium-permeable ion channels have been implicated in these processes, including TRPC channels.

Canonical transient receptor potential 6 (TRPC6) is a non-specific cation channel member of the transient receptor potential (TRP) superfamily of ion channels.[3–5] In addition to acting as a homo-tetramer, TRPC6 can form heteromeric channels with TRPC1, 3 and 7.[6, 7] TRPC6 is directly activated by diacylglycerol (DAG),[8] and is thought to act downstream of Gα_q_ coupled receptors, acting as a receptor-operated calcium effector.[9, 10] In light of its broad tissue and cell type expression, it is not surprising that TRPC6 is reported to influence multiple physiological and pathophysiological processes.[11–13] In humans, *TRPC6* mutations are a cause of autosomal dominant focal segmental glomerulosclerosis (FSGS); the majority of these act as gain-of-function.[14, 15] However, mice with a *Trpc6^E896K^* allele (corresponding to the human FSGS-associated, GOF E897K mutation) do not develop appreciable glomerular disease.[16] A mechanistic understanding of how TRPC6 mutations lead to kidney disease remains absent.

TRPC6 is abundantly expressed in platelets,[17–20] but the channel’s role in these cell fragments is still incompletely understood. *Trpc6* knockout does not affect most agonist stimulated platelet aggregation, integrin activation or degranulation, though there are conflicting data on a role for TRPC6 in response to thromboxane receptor activation.[18, 21] TRPC6, together with TRPC3, has been proposed to act as a co-incident detector of thrombin and collagen stimulation to induce phosphatidylserine (PS) exposure.[20] PS exposure generates a procoagulant surface that enhances thrombin generation.[22] Finally, TRPC6 is required for the exaggerated platelet activation and inflammatory response seen in platelet-specific CFTR knockout mice, but its loss does not affect wild-type platelet function.[23] In the current study, we compared platelets from wild-type, *Trpc6^E896K/E896K^*, and *Trpc6^DN/DN^* mice to further characterize the role of TRPC6 in platelet function. We found that GOF TRPC6 impacts several signaling pathways, including PS exposure, Erk1/2 and MLC2 phosphorylation, talin cleavage and clot retraction, but that most of these pathways are not affected by the loss of TRPC6 channel activity. These studies suggest that TRPC6 GOF mutants do not simply enhance wild-type TRPC6 responses, but can activate effector pathways not normally impacted by TRPC6 channel activity.

## Materials and methods

### Materials

Chemicals were purchased from Sigma Aldrich unless otherwise specified. GSK1702934A (GSK) was from Focus Biomolecules, TRPC6-PAM-C20 from Tocris, and U46619 from Cayman Chemical. Antibodies were from the following sources: Actin (#3700), Erk1/2 (#4695), phospho-Erk1/2 (#4370), phospho-MLC2 (#3675), Talin (#4021), all from Cell Signaling Technology; TRPC6 (ACC-017) from Alomone; FITC AnnexinV (556419) and PE CD41 (561850) from BD Biosciences.

### Mouse Studies

All animal procedures were approved by the Beth Israel Deaconess Medical Center (BIDMC) Animal Care and Use Committee, and carried out in accordance with the National Institutes of Health Guide for the Care and Use of Laboratory Animals. Animals carrying mutant *Trpc6* alleles, *Trpc6^E896K/E896K^* and *Trpc6^DN/DN^* (introducing an LFW to AAA pore-mutation), were generated at the BIDMC transgenic core using CRISPR/Cas9 with two site-specific guide RNAs (PNA Bio) per locus, Cas9 nickase (PNA Bio), and a single stranded DNA oligo (IDT) carrying the desired mutant sequence. Guide RNA and oligo DNA sequences are listed in Table 1 and 2, respectively. Genomic DNA from founder mice was amplified by PCR and screened by Sanger sequencing. Founders carrying the desired mutation were mated with FVB/NJ animals (Jackson Laboratory). After at least 5 backcrosses with FVB/NJ animals, heterozygous animals were mated to generate homozygous animals and wild-type littermates. *Trpc6^E896K/E896K^* and *Trpc6^DN/DN^* lines were then maintained by mating homozygous animals. Genotyping of the *Trpc6^E896K^* locus was performed using a custom TaqMan SNP assay; the *Trpc6^DN^* locus was genotyped using standard PCR reactions to detect the wild-type and DN alleles. *Trpc6^-/-^* mice, previously reported,[24] were obtained from Jackson Laboratory. After crossing the mice with C57BL/6J, heterozygous *Trpc6^+/-^* mice were crossed to generate littermate *Trpc6^-/-^* and *Trpc6^+/+^* animals.

**Table 1.**
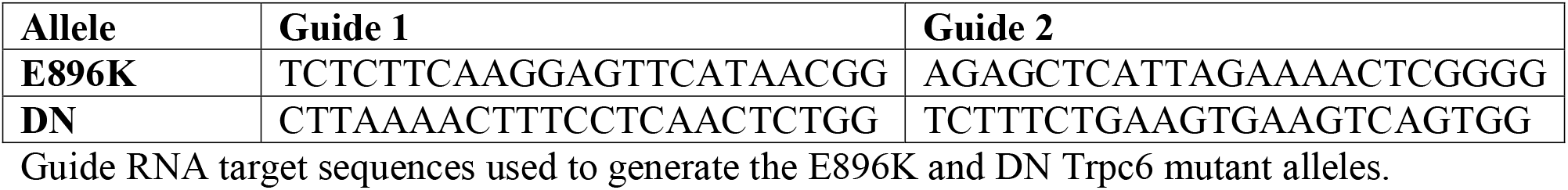
Guide RNA sequences.

**Table 2.**
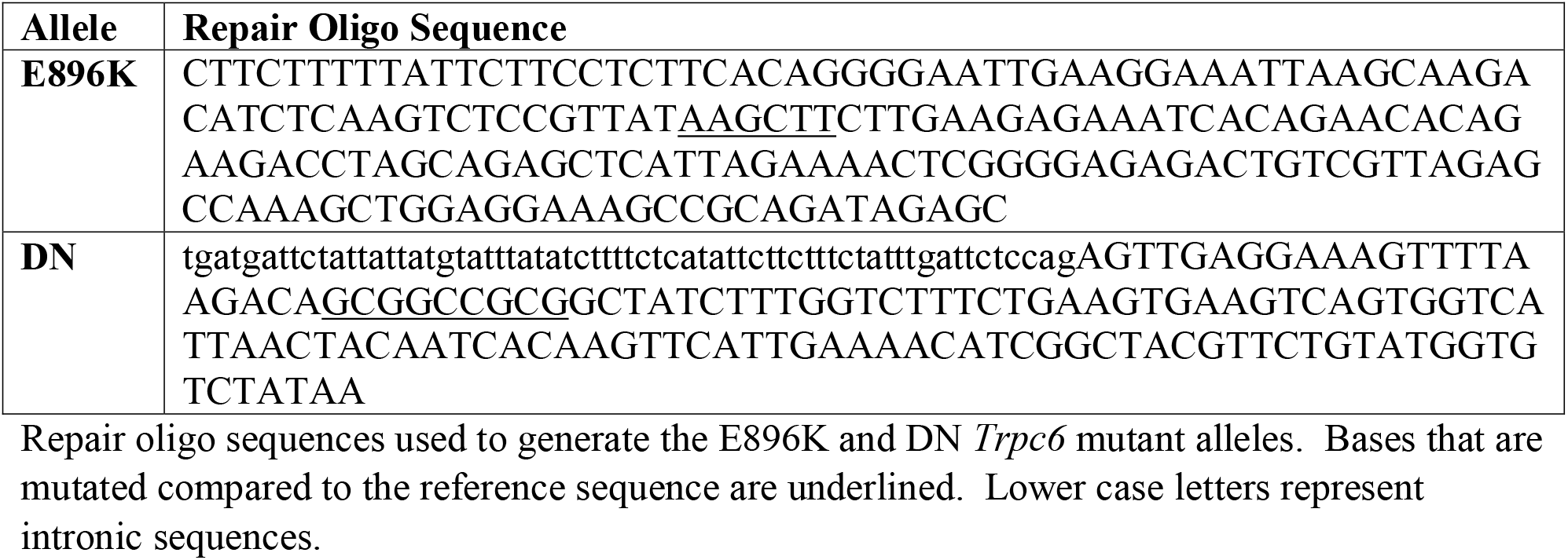
Repair oligo sequences.

Whole blood was obtained via cheek pouch bleeding and collected in EDTA-containing microtainers (BD 365974). Blood counts were analyzed on a Hemavet 950 FS at the BIDMC Small Animal Imaging core facility.

Tail bleeding time was assessed using standard protocols.[19] Briefly, animals were anesthetized with inhaled isoflurane and maintained on a warming blanket. The tail was severed 5 mm from the tip with a sterile scalpel and immediately placed in sterile PBS warmed to 37°C. The time taken until bleeding stopped, and did not resume for at least 60 seconds, was recorded. The experiment was terminated if bleeding persisted after 15 minutes.

### Platelet Isolation

Mouse platelets were isolated using modified standard procedures.[25] Whole blood was obtained by cardiac puncture from deeply anesthetized mice using 1ml syringes prefilled with 100 μl of 10% sodium citrate solution. Samples were diluted with an equal volume of HEPES-Tyrodes buffer without calcium (20 mM HEPES, 135 mM NaCl, 2.8 mM KCl, 1mM MgCl_2_, 12 mM NaHCO_3_, 0.4 mM NaH_2_PO_4_, 0.25% BSA, 5.5 mM glucose, pH 7.4) supplemented with 0.04 u/ml Apyrase and 2 μM PGE_1_, and an additional one-tenth volume of sodium citrate solution. Platelet rich plasma (PRP) was obtained after centrifugation at 220g for 10 minutes at room temperature. After resting at 37°C for 15 minutes, 1:6 volume ACD solution (75 mM Trisodium citrate, 42 mM citric acid, 136 mM glucose) was added and platelets pelleted at 500g for 10 minutes. Platelets were washed once in 500 μl HEPES-Tyrode buffer with apyrase and PGE_1_, before being resuspended in HEPES-Tyrodes buffer with or without CaCl_2_ for use in downstream experiments.

### Calcium Imaging

Intracellular calcium levels were assessed using Fura-2 fluorescence ratio measurement (Fura-2 QBT kit R8197, Molecular Devices) on a FlexStation III reader with automated pipetting at the Harvard ICCB-Longwood screening facility. Platelets were resuspended in HEPES-Tyrodes buffer with Fura-2 and 1.2 mM CaCl_2_; 100 μl of platelet suspension were added to each well of a 96-well plate. Platelets were incubated with Fura-2 for 1hr at 37°C before beginning the assay, and were maintained at 37°C in the FlexStation III reader throughout the experiment. Samples were excited at wavelengths of 340 and 380 nm every 5 s for a total of 330 s, with emissions measured at 510 nm. Baseline fluorescence measurements were obtained for 30 seconds before stimulation by the addition of agonist in 50 μl HEPES-Tyrodes buffer with 1.2 mM CaCl_2_. Final concentrations of agonists/inhibitors were as follows: 10 μM ADP, 3 μM CFTR-inhibitor 172, 10 μM fluoxetine, 50 μM GSK1702934A, 0.5 u/ml thrombin, 10 μM TRPC6-PAM-C20, 5 μM U46619, 10 μM Y-27632. In experiments examining the effects of pre-incubation with inhibitors, drug was added at the start of the Fura-2 incubation.

### Flow Cytometry

For phosphatidylserine exposure experiments, platelets were resuspended in HEPES-Tyrode buffer supplemented with 1 mM CaCl_2_, mixed with an equal volume of buffer containing agonist or carrier, and incubated at room temperature for 10 minutes. Agonists and inhibitors were used at the same concentration as for Fura2 imaging experiments, above. 10 μl of platelet suspension was mixed with 40 μl of staining solution (HEPES-Tyrode buffer supplemented with 2.5 mM CaCl2, 1:50 PE rat anti-mouse CD41 antibody (BD Biosciences 561850), and 3:50 FITC Annexin V (BD Biosciences 556419)). Control staining reactions containing one or no fluorophore were performed in parallel. After a 10 minute room-temperature incubation, platelets were diluted with 200 μl of 1.25% paraformaldehyde solution. Cell-surface P-selectin exposure and integrin αIIbβ3 activation was assayed using a two-color mouse platelet activation kit (Emfret Analytics D200) following the supplied protocol. Flow cytometry was performed at the BIDMC flow cytometry core facility on a Beckman Coulter CytoFLEX LX.

### Western Blotting

Washed platelets were resuspended in HEPES-Tyrode buffer supplemented with 1 mM CaCl_2_, mixed with agonists or inhibitors as indicated, and incubated at room temperature. At the indicated time points, aliquots of platelet suspension were mixed with 4x sample loading buffer containing β-mercaptoethanol and immediately incubated at 95°C for 5 minutes. SDS-PAGE and Western blotting were performed as previously described [26]. Antibodies against the following antigens were utilized: actin (CST #3700, 1:1000), Erk1/2 (CST #4695, 1:1000), phospho-Erk1/2 (CST #4370, 1:2000), phospho-MLC2 Ser19 (CST #3675, 1:1000), talin (CST #4021; 1:1000), TRPC6 (Alomone ACC-017, 1:500).

### Clot Retraction

Clot retraction was assayed following standard protocols.[27] Citrated PRP was collected as outlined above. 100 μl of PRP and 5 μl of pelleted RBCs (to provide clot contrast) were diluted to 500 μl final volume with HEPES-Tyrode buffer (final 1.5 mM CaCl_2_ concentration), supplemented with or without 1 u/ml thrombin, and incubated at room temperature. Tubes were imaged every 15 or 30 minutes for two hours, and two dimensional clot area calculated as a percentage of the total fluid area at each timepoint. A clot “area under the curve” was calculated by summing the clot area for each sample at each timepoint.

### Statistical Analysis

All statistical analyses were performed using GraphPad Prism version 9 or later. The statistical tests utilized for each experiment are specified within the corresponding figure legends. Symbols use for pair-wise comparison adjusted p-values are: *, p<0.05; **, p<0.01; ***, p<0.001; ****, p<0.0001.

## Results

### TRPC6 platelet expression and function in *Trpc6* mutant animals

Utilizing CRISPR/Cas9, we generated two mouse lines carrying mutations in the *Trpc6* gene. The *Trpc6^E896K^* line (also termed KI) encodes an E896K amino acid change in the TRPC6 protein to mimic the human FSGS disease associated mutation E897K.[14] The human mutation demonstrates gain-of-function properties.[14, 26, 28] A second mutation, *Trpc6^DN^* (also termed DN) encoding for an LFW to AAA amino acid change in the pore domain of TRPC6, acts as a dominant negative mutant.[6] Homozygous *Trpc6^E896K/E896K^* and *Trpc6^DN/DN^* mice are viable, fertile, and showed no overt phenotype. Furthermore, *Trpc6^E896K/E896K^* animals do not develop albuminuria or histological evidence of kidney disease through 6 months of age.[16]

As TRPC6 is abundant in platelets,[17–20] we compared TRPC6 levels in platelets from our mouse lines. Specificity of the anti-TRPC6 antibody was confirmed using platelet lysates from *Trpc6^-/-^* animals (Fig 1A). TRPC6 protein was detected in wild-type, *Trpc6^E896K/E896K^* and *Trpc6^DN/DN^* platelets (Fig 1B). TRPC6 abundance was slightly lower in KI platelets. Variability in the different molecular weight forms of TRPC6 between genotypes was similar to our previous observations in overexpression systems.[28] Platelet counts were similar across all three genotypes, as were other aspects of a complete blood count (Table 3).

**Fig 1.**
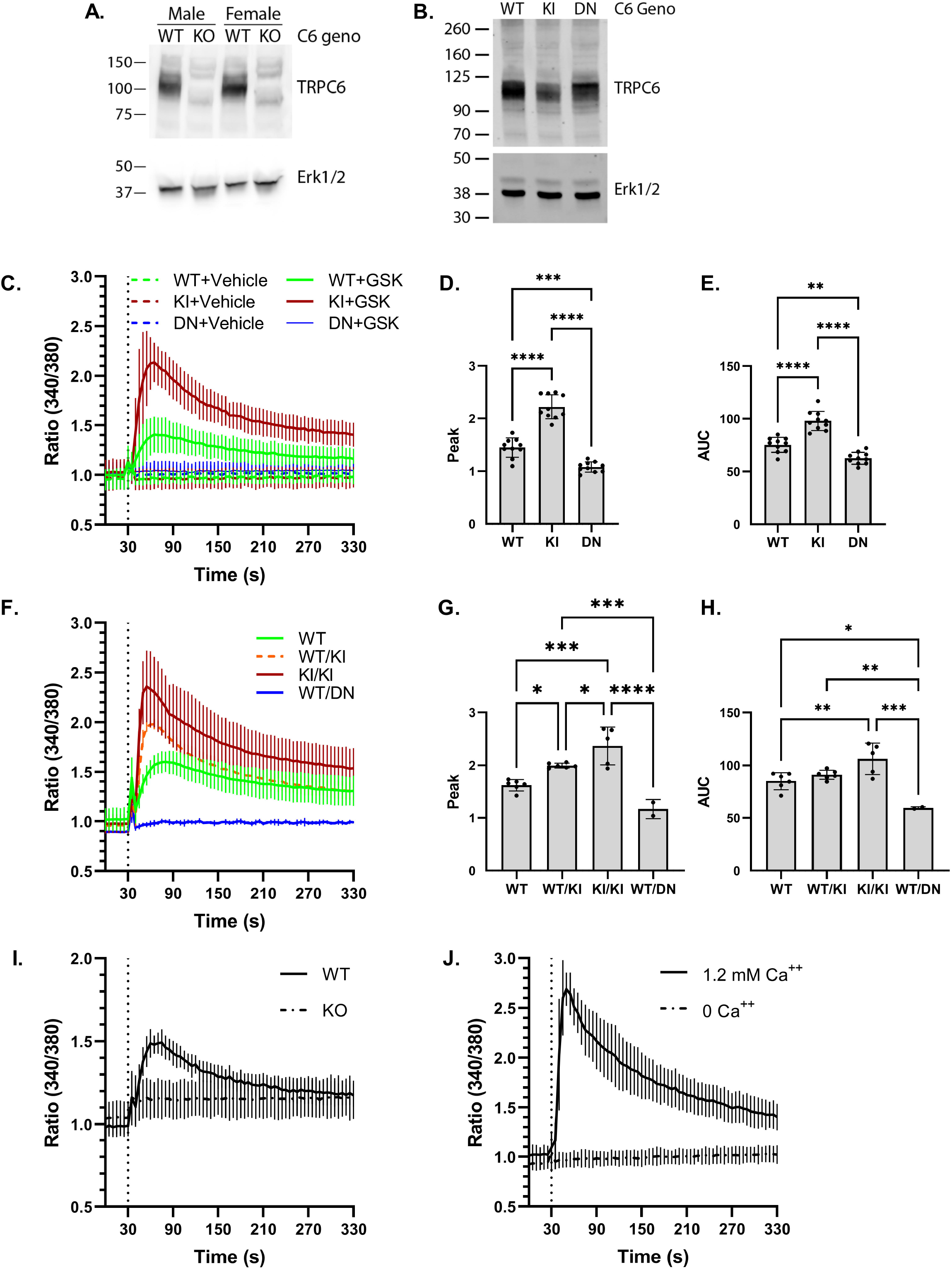
Characterization of TRPC6 expression and GSK1702934A response in *Trpc6* mutant platelets. A, platelet lysates from wild-type (WT) and *Trpc6^-/-^* (KO) littermate animals were analyzed by SDS-PAGE and western blot. Anti-TRPC6 antibodies (top) detected a major band around 105 kD in WT but not KO animals. Erk1/2 blot (bottom) is shown as a loading control. B, platelet lysates obtained from wild-type (WT), *Trpc6^E896K/E896K^* (KI), and *Trpc6^DN/DN^* (DN) animals were subject to western blot using antibodies against TRPC6, and Erk1/2. C, timecourse of Fura-2 fluorescence ratio (340/380) in WT, KI and DN platelets. Platelets were treated with GSK (final concentration 50 μM) or DMSO vehicle after 30 seconds (dashed vertical line). Shown are mean ± SD; n=8-10 using platelets from 4-6 animals per genotype. D, peak poststimulation Fura-2 fluorescence ratios from GSK-stimulated traces shown in (C); shown are the mean ± SD, and values from individual experiments. One-way ANOVA with Tukey’s multiple comparisons test. E, area under the curve (AUC, arbitrary units) of the Fura-2 fluorescence ratio stimulation was calculated for each experiment shown in C. Shown are the mean and values from individual experiments. One-way ANOVA with Tukey’s multiple comparisons test. F, Fura-2 time-course in GSK-stimulated platelets from WT, *Trpc6^+/E896K^* (WT/KI), *Trpc6^E896K/E896K^* (KI/KI), and *Trpc6^+/DN^* (WT/DN) platelets. Shown are mean ± SD; n= 2-6 animals/genotype. Peak Fura-2 fluorescence ratio (G), and area under the curve (H), over baseline after GSK stimulation are shown for platelets from the indicated *Trpc6* genotype. One-way ANOVA with Tukey’s multiple comparisons test. I, time course of Fura-2 fluorescence ratio in platelets from wild-type and *Trpc6^-/-^* (KO) animals stimulated with GSK. Data are mean ± SD; n=8/group using platelets from 2 animals per genotype. J, Fura-2 time course in *Trpc6^E896K/E896K^* platelets stimulated with GSK in buffer with or without extracellular calcium. Data are mean ± SD; n=3/condition using platelets from 2 animals.

**Table 3.** Blood Count Values by *Trpc6* Genotype.

|  | Wild-type | E896K/E896K | DN/DN |
| --- | --- | --- | --- |
| WBC (K/μL) | 3.24 ± 1.11 | 3.14 ± 0.55 | 4.52 ± 1.13 |
| Hgb (g/dL) | 12.8 ± 1.2 | 12.9 ± 1.7 | 13.2 ± 0.8 |
| Platelets (K/μL) | 736 ± 334 | 516 ± 187 | 620 ± 346 |
| **MCV (fL)** | 45.7 ± 0.9 | 45.9 ± 0.9 | 45.8 ± 0.7 |
| **MPV (fL)** | 4.92 ± 0.38 | 4.98 ± 0.36 | 5.06 ± 0.36 |
| **Neutrophils (K/μL)** | 0.84 ± 0.24 | 0.52 ± 0.15 | 1.01 ± 0.43 |
| **Lymphocytes (K/μL)** | 2.51 ± 0.83 | 2.45 ± 0.38 | 3.24 ± 0.82 |
| **Monocytes (K/μL)** | 0.10 ± .04 | 0.12 ± 0.03 | 0.16 ± 0.15 |

Values shown are means ± standard deviation. None of the parameters show statistically significant differences between genotypes by one-way ANOVA; n=5 per group.

Fura-2-based ratiometric calcium fluorimetry was performed to assess the functional consequences of the E896K and DN mutations on TRPC6 channel function. Treating wild-type platelets with GSK1702934A (GSK) induced a rapid, but transient rise in the F340/F380 ratio (Fig 1C). *Trpc6^DN/DN^* platelets demonstrated no significant response, while *Trpc6^E896K/E896K^* platelets demonstrated a higher peak and a prolonged elevation in the fluorescence ratio (Fig 1C-E). *Trpc6^+/DN^* platelets also failed to respond to GSK, consistent with the mutation’s dominant negative phenotype (Fig 1F). *Trpc6^+/E896K^* platelets showed an intermediate Fura-2 response relative to wild-type and *Trpc6^E896K/E896K^* platelets (Fig 1F-H). Platelets from *Trpc6^-/-^* animals failed to respond to GSK stimulation, suggesting that TRPC6 is the only major target of this agonist that influences calcium influx in platelets (Fig 1I). Omitting extracellular calcium completely abolished the agonist effect in *Trpc6^E896K/E896K^* platelets (Fig 1J). In sum, these results suggest that TRPC6 is required for extracellular calcium influx in response to GSK in platelets, and that the E896K and DN mutant channels display properties similar to what has been reported in *in vitro* overexpression studies.[6, 14]

Activation of Gα_q_ coupled receptors is known to activate TRPC6 in heterologous overexpression systems.[8–10] We therefore examined calcium signaling downstream of thrombin, ADP and the thromboxane A_2_ receptor agonist, U46619, in *Trpc6* mutant platelets. No differences were noted in Fura-2 fluorimetry between wild-type, *Trpc6^E896K/E896K^* and *Trpc6^DN/DN^* platelets in response to ADP (Fig 2A-C). While the peak F340/380 values after thrombin stimulation similarly did not differ between genotypes, the area under the curve was modestly higher in the *Trpc6^E896K/E896K^* platelets compared to *Trpc6^DN/DN^* platelets, but not significantly different between wild-type and *Trpc6^DN/DN^* samples (Fig 2D-F). Responses to U46619 were similar between *Trpc6^E896K/E896K^* and *Trpc6^DN/DN^* platelets (Fig 2G-I). Given this result, wild-type platelets were not tested in keeping with the three R’s tenet in animal research. Together, these results suggest that TRPC6 may play a modest role in calcium signaling downstream of thrombin, but is not involved downstream of ADP or U46619.

**Fig 2.**
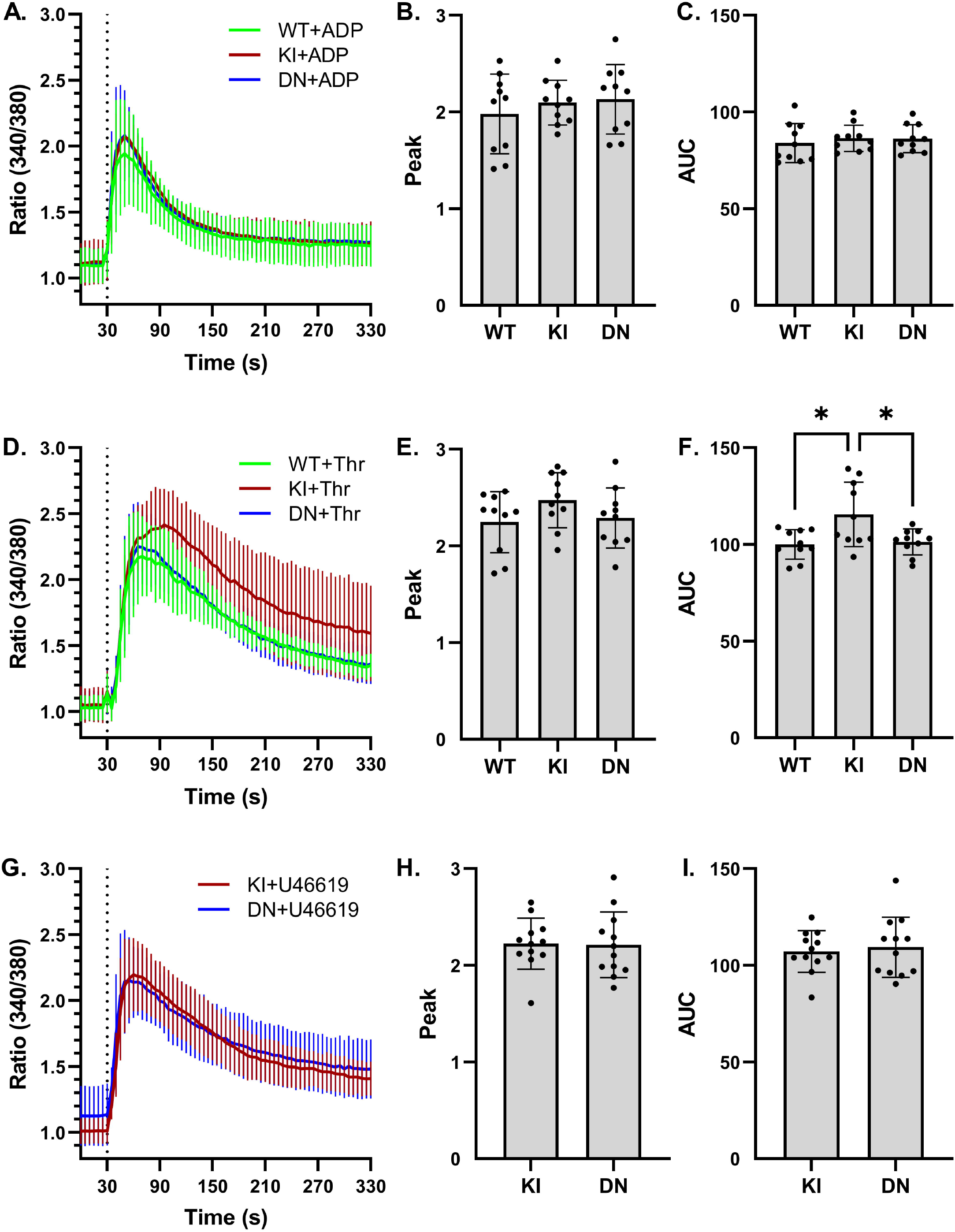
Effect of *Trpc6* genotype on ADP, thrombin, and U46619-stimulated platelet calcium influx. A, Fura-2 fluorescence ratio (340/380) time-course in wild-type (WT), *Trpc6^E896K/E896K^* (KI), and *Trpc6^DN/DN^* (DN) platelets stimulated with ADP (final concentration 10 μM) after 30 seconds (dashed vertical line). Shown are mean ± SD; n=8-10 using platelets from 6 animals per genotype. B, peak post-stimulation Fura-2 fluorescence ratios from ADP-stimulated traces shown in (A); shown are the mean and values from individual experiments. C, area under the curve (AUC, arbitrary units) of the Fura-2 fluorescence ratio was calculated for each experiment shown in A. Shown are the mean and values from individual experiments. D, Fura-2 time course as in (A) above, but with thrombin (final concentration 0.5 u/ml) added at 30 seconds; n=8-10 using platelets from 6 animals per genotype. Peak Fura-2 fluorescence ratio (E), and area under the curve (F), after thrombin stimulation of platelets from the indicated *Trpc6* genotype. G, Fura-2 time course in KI and DN platelets stimulated with 5 μM U46619; n=12 using platelets from 5 animals per genotype. Peak fluorescence ratio (H) and AUC (I) after stimulation did not differ between genotypes. One-way ANOVA with Tukey’s multiple comparisons test. Only statistically significant pairwise comparisons are shown.

### Effects of *Trpc6* mutations on integrin αIIbβ3 activation and P-selectin exposure

Activation of integrin αIIbβ3, and cell surface exposure of P-selectin via degranulation, are two common responses of platelets to agonist stimulation. Previous studies report that *Trpc6* deficiency does not affect these responses to various stimuli, including ADP and thrombin, though there is disagreement as to whether thromboxane A2 receptor agonist response is blunted.[18, 21] The effect of *Trpc6* genotype on integrin αIIbβ3 activation and P-selectin surface expression was assessed by flow cytometry (Fig 3A-B). In line with the lack of effect on ADP and U46619 induced calcium transients (Fig 2), *Trpc6* genotype did not affect αIIbβ3 activation or P-selectin surface expression in response to these stimuli (Fig 3A-B). There was also no difference in response to thrombin-mediated activation. GSK alone did not activate wild-type or *Trpc6^DN/DN^* platelets, but did induce low level activation of *Trpc6^E896K/E896K^* platelets. Combined stimulation with ADP and GSK enhanced P-selectin exposure in wild-type and *Trpc6^E896K/E896K^* platelets compared to *Trpc6^DN/DN^* platelets. Taken together, these results suggest that the increased calcium influx mediated by GOF TRPC6 is sufficient to induce platelet activation, while that induced by wild-type channel can augment activation mediated by the weak agonist ADP, but is insufficient on its own.

**Fig 3.**
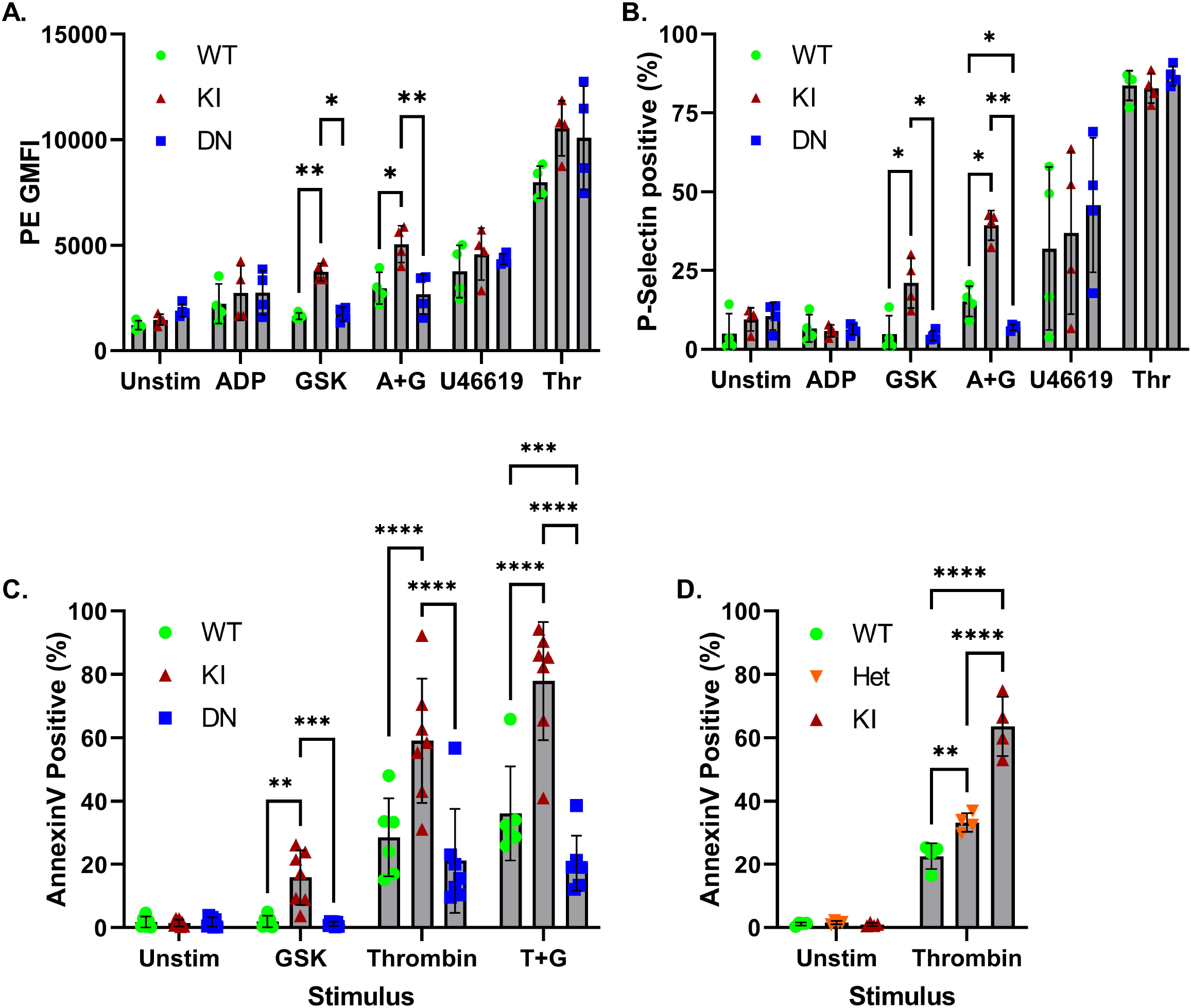
Differential effects of *Trpc6* genotype on agonist-induced integrin αIIbβ3 activation, P-selectin surface expression, and phosphatidylserine exposure. A-B, washed platelets from WT, KI, and DN animals were stimulated with the indicated agonists (A+G, ADP plus GSK; Thr, thrombin) in the presence of PE-conjugated anti-JON/A antibody (targeting activated integrin αIIbβ3) and FITC-conjugated antibody directed at P-selectin, fixed, and analyzed by flow cytometry. Geometric mean PE fluorescence intensity (A), and percentage of platelets with positive FITC labeling (B), are shown as means and individual values; n=4 animals per genotype. Two-way ANOVA with Tukey’s multiple comparisons test. Only statistically significant pairwise comparisons between genotypes subject to the same stimulus are shown. CD, washed platelets from the indicated *Trpc6* animals (Het, *Trpc6^+/E896K^*) were treated with the indicated stimuli, stained with anti-CD41 antibody and Annexin V, and subject to flow cytometry. The percentage of CD41 positive platelets staining for Annexin V is shown. C, percentage of Annexin V stained platelets from three *Trpc6* genotypes left unstimulated (unstim), treated with GSK, thrombin, or thrombin plus GSK (T+G); n=6-7 animals per genotype. Mixed-effects analysis with Tukey’s multiple comparisons test. D, Annexin V staining of platelets from wild-type, heterozygous and homozygous *Trpc6^E896K^* knock-in animals at baseline and after thrombin stimulation; n=4 per group. Two way ANOVA with Sidak’s multiple comparisons test. Only statistically significant pairwise comparisons are shown.

### *Trpc6* mutations affect phosphatidylserine exposure on platelets

In response to strong agonist signals, platelets expose phosphatidylserine on their plasma membrane, creating a highly pro-coagulant platelet surface.[22] TRPC3 and TRPC6 have been implicated in this process in response to dual activation by thrombin and CRP.[20] We therefore examined Annexin V staining by flow cytometry to assay phosphatidylserine exposure of platelets from wild-type, *Trpc6^E896K/E896K^* and *Trpc6^DN/DN^* mice (Fig 3C). At baseline, few platelets stained with Annexin V. GSK stimulation induced a small, but significant, increase in the percentage of Annexin positive platelets only in *Trpc6^E896K/E896K^* platelets. Thrombin stimulation induced a significant number of platelets from all genotypes to stain for Annexin, but the percentage was higher for the *Trpc6^E896K/E896K^* genotype. While wild-type and *Trpc6^DN/DN^* platelet responses to thrombin alone were similar to each other, the percentage of Annexin positive platelets after combined stimulation with thrombin and GSK was higher in wild-type platelets compared to *Trpc6^DN/DN^* platelets. In our hands, type I collagen or convulxin did not induce PS exposure, or enhance the effect of thrombin on PS exposure (data not shown). As a whole, these results suggest that TRPC6 channel activity is not necessary for PS exposure in response to thrombin, but that *Trpc6* gain-of-function mutations may enhance this process. Indeed, a higher percentage of *Trpc6^+/E896K^* platelets bound Annexin after thrombin stimulation than did wild-type platelets (Fig 3D).

### TRPC6 differentially modulates Erk and MLC2 phosphorylation

Both Erk1/2,[29] and myosin light chain 2 (MLC2),[30] phosphorylation are known downstream responses to calcium signaling in platelets, and of TRPC6 activation in other cell types.[26, 31–34] Therefore, we examined if TRPC6 activation, and genotype, affects these signaling pathways in platelets. Time course experiments in wild-type and *Trpc6^E896K/E896K^* platelets demonstrated peak Erk phosphorylation 5 minutes after GSK addition (Fig 4A). Wild-type platelets demonstrated a small, but consistent, increase in P-Erk1/2 levels 5 minutes after stimulation with GSK (Fig 4B, C). This response was absent in *Trpc6^DN/DN^* platelets, but augmented in *Trpc6^E896K/E896K^* platelets. GSK-mediated Erk activation was dependent on the presence of extracellular calcium (Fig 4D). Erk activation therefore appears to be a downstream target of TRPC6 activation in platelets. These results confirm our previous studies in heterologous overexpression systems that TRPC6 gain-of-function mutants enhance activation of the Erk1/2 signaling pathway.[26]

**Fig 4.**
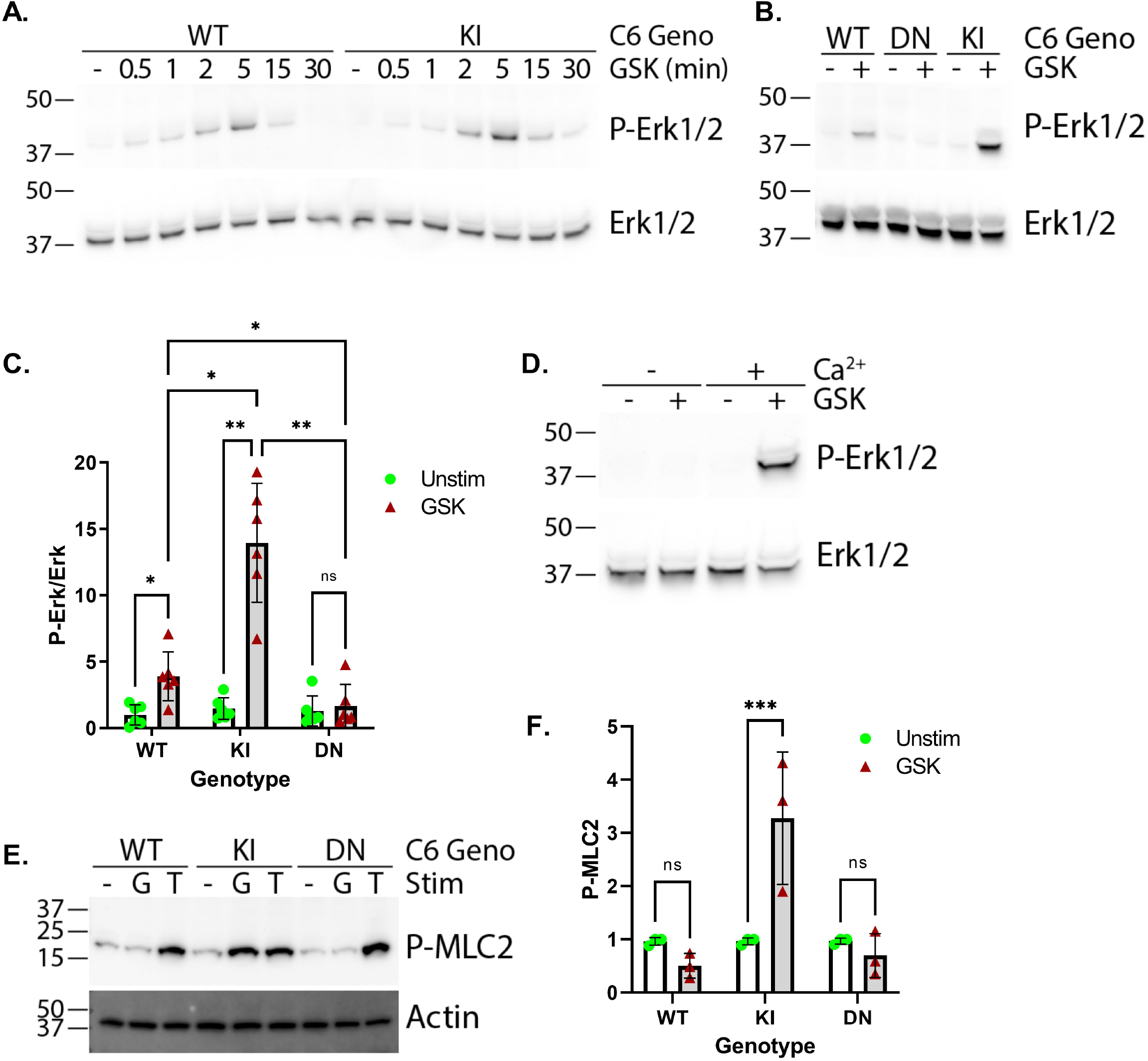
*Trpc6* mutations differentially affect Erk1/2 and MLC2 phosphorylation in platelets. Western blot analysis of platelet lysates from various *Trpc6* mice stimulated with or without GSK as indicated. A, time-course of phospho-Erk1/2 levels in WT and KI platelets after stimulation with GSK for the indicated time. Total Erk1/2 levels were utilized as loading controls (bottom panel). B, western blot for phospho-Erk1/2 (top) and total Erk (bottom) in platelet lysates from the indicated *Trpc6* genotype, treated with GSK or vehicle for 5 minutes prior to lysis. C, quantification of P-Erk1/2 to Erk ratios from western blot analysis; n=6 per group. Two-way ANOVA with Sidak’s multiple comparisons test. D, western blot of KI platelets treated with or without GSK in the absence or presence of extracellular calcium. E, phospho-MLC2 (Ser19) blot of platelets unstimulated (-), or stimulated with GSK (G) or thrombin (T) for 5 minutes. Actin served as a loading control. F, quantification of phospho-MLC2 (Ser19) levels after 5 minutes of GSK stimulation; n=3. Two-way ANOVA with Sidak’s multiple comparisons test.

In contrast to its effect on Erk1/2 phosphorylation, GSK increased phospho-MLC2 levels in *Trpc6^E896K/E896K^*, but not in wild-type or *Trpc6^DN/DN^*, platelets (Fig 4E, F). Thrombin treatment induced robust increases in MLC2 phosphorylation regardless of *Trpc6* genotype. These results suggest that MLC2 phosphorylation is not normally a downstream response to TRPC6 activation in platelets, but can be induced by activation of GOF TRPC6.

### Gain-of-function TRPC6 activation induces talin cleavage and delays clot retraction

Phosphatidylserine-exposed platelets tend to inactivate integrin αIIbβ3, in part through the calcium-dependent cleavage of the focal adhesion protein, talin.[35, 36] We found that GSK stimulation of *Trpc6^E896K/E896K^* platelets, but not wild-type or *Trpc6^DN/DN^* platelets, induced talin cleavage as assessed by western blot (Fig 5A). This response was seen within a minute after stimulation of *Trpc6^E896K/E896K^* platelets, but remained absent even after 30 minutes of stimulation of wild-type platelets (Fig 5B).

**Fig 5.**
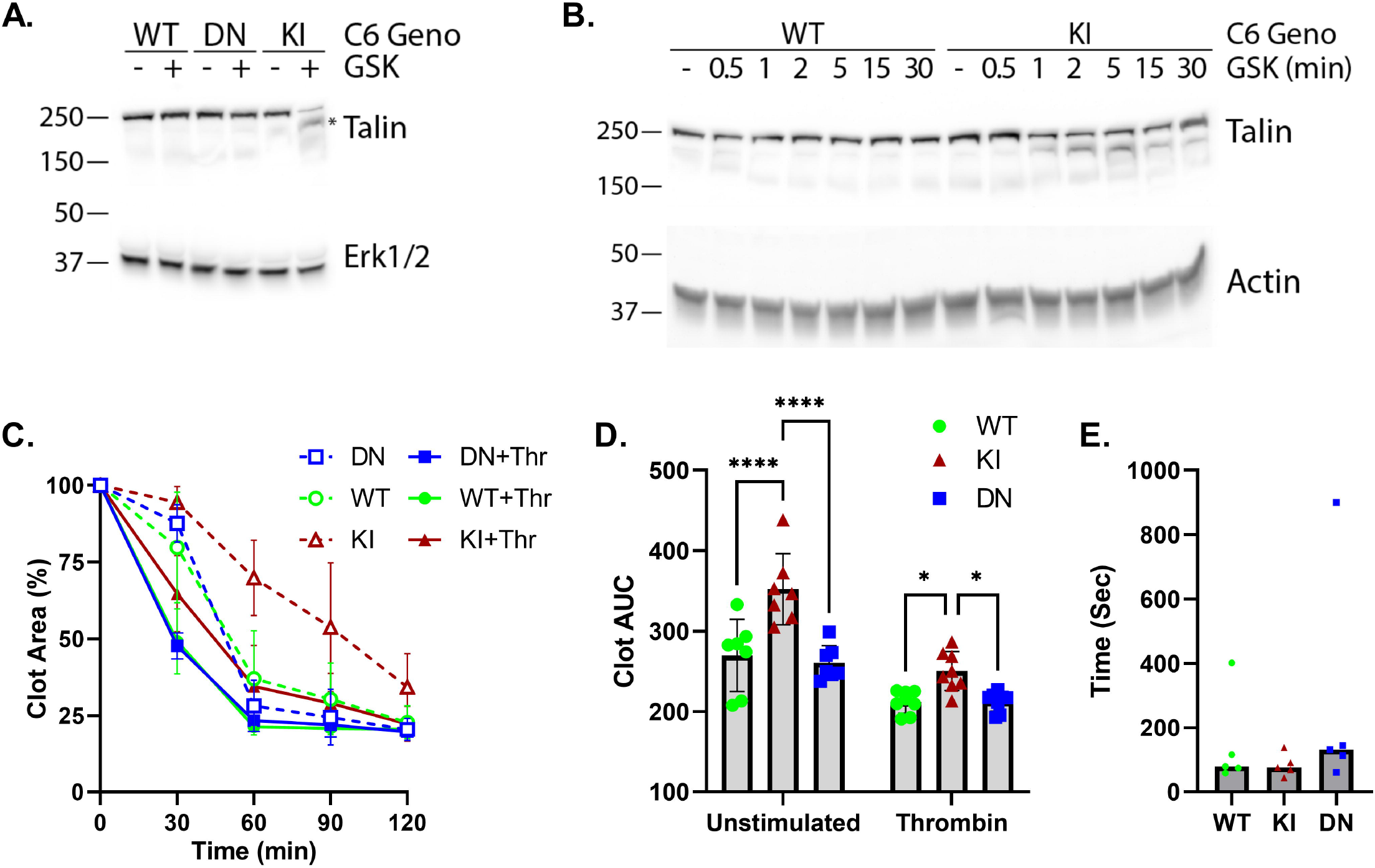
Effect of *Trpc6* genotype on talin cleavage, clot retraction and tail bleeding times. A, western blot analysis of platelet lysates of the indicated *Trpc6* genotype treated with or without GSK for 5 minutes. Lysates were probed for Talin (top) and Erk1/2 (bottom). The * points out the lower molecular weight Talin band corresponding to cleaved protein. B, time-course of Talin cleavage. WT and KI platelets were stimulated with GSK for the indicated time prior to lysis and western blot analysis. Actin served as a loading control. C-D, platelet-rich plasma (PRP) from WT, KI, and DN animals was incubated in the presence of extracellular calcium without or with thrombin stimulation. Clot area was calculated as a percentage of total sample area at various times. C, time course of clot area. Shown are means ± SD, n= 7-8 per group. D, clot area under the cure (AUC) was calculated by summing the clot areas across all time-points for each sample. Shown are means ± SD, and individual values. Mixed-effects analysis with Tukey’s multiple comparisons test. E, tail bleeding times in animals with WT, KI and DN *Trpc6* genotypes. Shown are median and individual values; n=5 per group.

Cleavage of platelet focal adhesion proteins, including talin, has been associated with relaxed fibrin clot retraction.[36] There are conflicting reports on the effect of *Trpc6* knockout on clot retraction and tail bleeding time.[18, 19, 21] In our studies, PRP from *Trpc6^E896K/E896K^* animals demonstrated delayed clot retraction compared to PRP from other genotypes, both in the absence of exogenous stimulus, and after thrombin addition (Fig 5C, D). No differences were seen between wild-type and *Trpc6^DN/DN^* genotypes. In contrast to the clot retraction kinetics, tail bleeding time did not appreciably differ between *Trpc6* genotypes (Fig 5E).

### Pharmacological modulation of TRPC6-mediated calcium influx and platelet activation

A positive allosteric modulator of TRPC6, TRPC6-PAM-C20 (C20), has been reported to enhance the response of TRPC6 to stimulation by OAG or GSK in human platelets.[37] We examined whether C20 has a similar effect on murine TRPC6, and whether it might further enhance mutant TRPC6^E896K^ activity. In wild-type platelets, C20 enhanced GSK-induced calcium influx, but had no effect on its own (Fig 6A-C). This effect was also seen in *Trpc6^E896K/E896K^* platelets, arguing that the positive allosteric effect of C20 acts via a different, and additive, mechanism to that of the gain-of-function E896K mutation.

**Fig 6.**
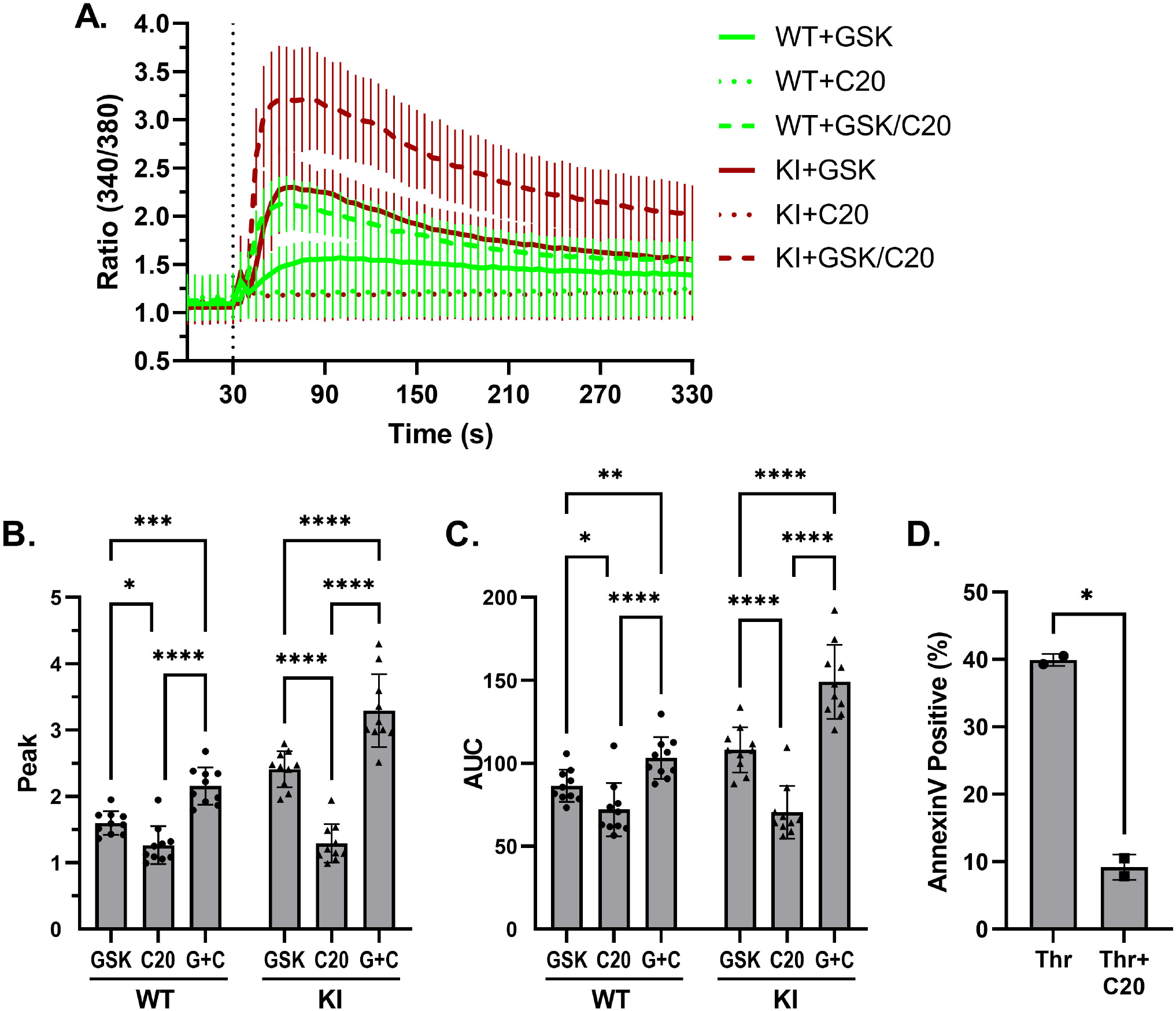
TRPC6-PAM-C20 enhances wild-type and mutant TRPC6-mediated calcium influx. A, Fura-2 fluorescence ratio (340/380) time-course in WT and KI platelets stimulated with the indicated combinations of C20 (10 μM) and GSK (50 μM) after 30 seconds. Shown are mean ± SD; n=10 using platelets from 5 animals per genotype. The post-stimulation Fura-2 fluorescence ratio peak (B), and area under the curve (C) are shown as means and individual values. Two way ANOVA with Tukey’s multiple comparisons test. D, percentage of DN platelets staining with Annexin V after being stimulated with thrombin (Thr) or thrombin plus C20; n=2 per group. Paired t-test.

We hoped to examine how C20 affects TRPC6-dependent phosphatidylserine exposure next. However, C20 inhibits thrombin-induced Annexin V staining in *Trpc6^DN/DN^* platelets (Fig 6D), suggesting it has TRPC6-independent effects, and preventing interpretation of its effects on TRPC6-dependent PS exposure.

Inhibitors of acid sphingomyelinases, including fluoxetine, are reported to inhibit TRPC6 channel activity [38–41]. We confirmed that pre-incubation of platelets with fluoxetine inhibits GSK induced calcium influx in wild-type and *Trpc6^E896K/E896K^* platelets (Fig 7A). In contrast, exposure to fluoxetine did not alter ADP-induced calcium transients (Fig 7B). The action of fluoxetine is likely indirect, as addition of fluoxetine one minute before GSK had a minimal effect on Fura2 fluorescence, as compared to a one hour pre-incubation (Fig 7C-E). Inhibition of GSK, but not ADP, stimulation by fluoxetine was mimicked by sertraline, but not by citalopram or venlafaxine (Fig 7 F, G), in line with their reported relative activities as functional inhibitors of acid sphingomyelinases, as opposed to their ability to inhibit serotonin uptake [41]. The effect of fluoxetine extended to Annexin V staining of *Trpc6^E896K/E896K^* platelets (Fig 7H). Preincubation completely abolished the increase in Annexin V staining upon GSK stimulation. Fluoxetine also decreased the percentage of Annexin V positive platelets after thrombin or thrombin plus GSK stimulation, to a similar level. These results suggest the presence of both fluoxetine sensitive, and insensitive, mechanisms of PS exposure downstream of thrombin stimulation.

**Fig 7.**
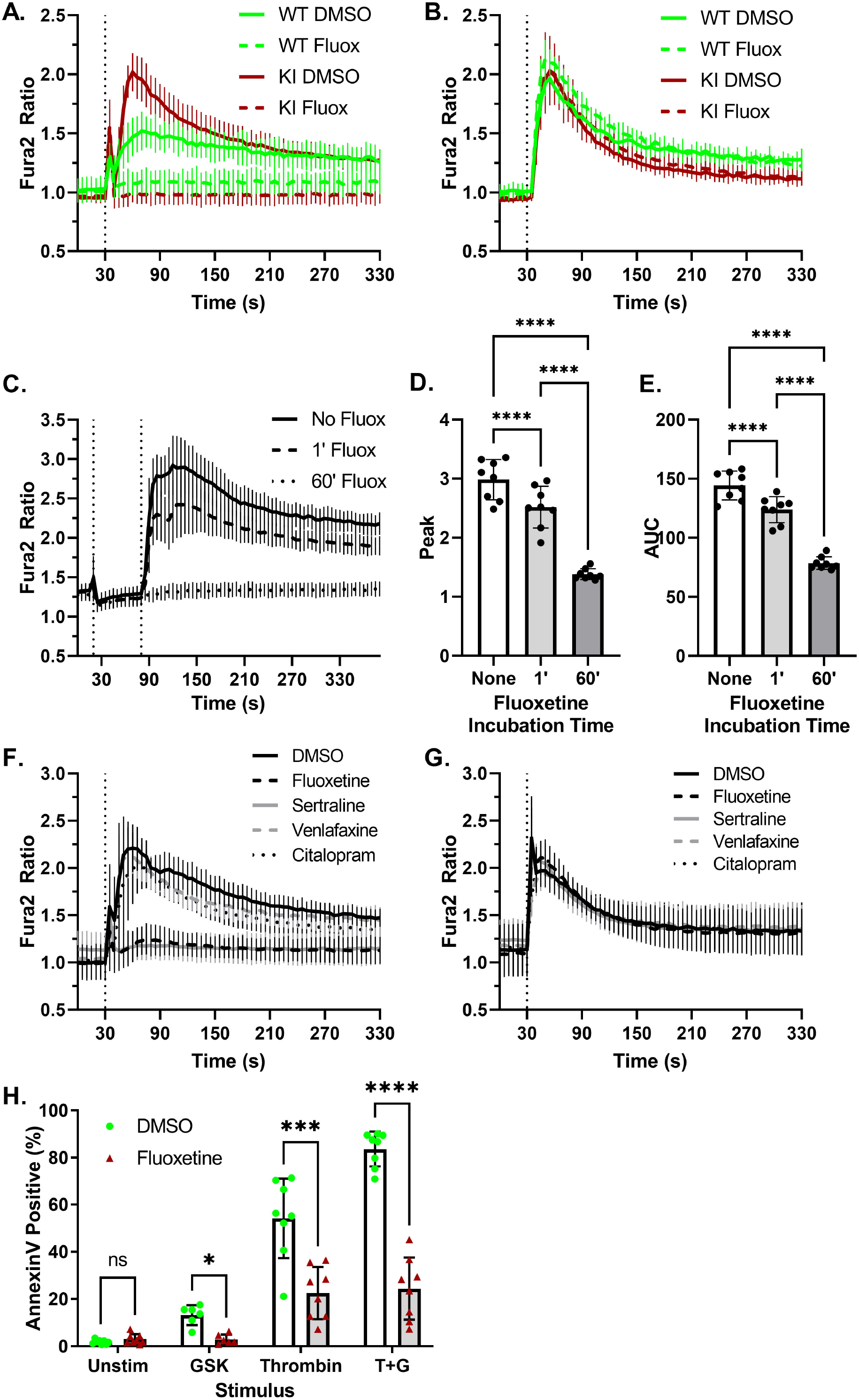
Fluoxetine inhibits TRPC6-mediated calcium influx and phosphatidylserine exposure. A, WT and KI platelets, pre-incubated in the presence of vehicle (DMSO) or fluoxetine (10 μM), were subject to Fura-2 ratiometric imaging. GSK was added after 30 seconds. Shown are mean ± SD, n=3-6. B, platelets were pretreated as in (A) followed by stimulation with ADP. Shown are mean ± SD, n=2-4. C, KI platelets were pre-incubated with (60’ Fluox) or without fluoxetine followed by Fura2 analysis. Fluoxetine (1’ Fluox) or vehicle (No Fluox and 60’ Fluox) was added to the platelets after 20 seconds (first vertical line) followed by GSK at 80 seconds; mean ± SD, n=8. Fura-2 fluorescence ratio peak (D) and area under the cure (E) are shown as mean and individual values. One way ANOVA with Tukey’s multiple comparisons test. F-G, Fura-2 imaging of KI platelets, pre-incubated with DMSO or the indicated drug (all 10 μM), and stimulated with GSK (F) or ADP (G) after 30 seconds. Shown are mean ± SD, n=5-6. H, percentage of *Trpc6^E896K/E896K^* platelets staining for Annexin V, after pre-incubation with carrier (DMSO) or fluoxetine, followed by stimulation with vehicle (Unstim), GSK, thrombin, or thrombin plus GSK (T+G); n=6-8 animals per group. Mixed effects analysis with Sidak’s multiple comparisons test.

Loss of CFTR is reported to enhance agonist-mediated platelet activation in a TRPC6-dependent manner [23], while genetic or pharmacologic interference of Rho-associated coiled-coil serine/threonine kinase 1 (ROCK1) enhances platelet PS exposure [42]. We therefore examined whether pharmacological inhibition of CFTR and ROCK affects TRPC6-mediated platelet activities (Fig 8). CFTR-inhibitor 172 did not significantly alter GSK-mediated calcium transients in either wild-type or *Trpc6^E896K/E896K^* platelets (Fig 8A-C). The inhibitor also had no effect on Annexin V staining regardless of genotype or stimulus (Fig 8G). The ROCK inhibitor, Y-27632, had no appreciable effect on calcium transients in wild-type platelets, but did slightly suppress the peak Fura-2 ratio, but not AUC, in GSK-stimulated *Trpc6^E896K/E896K^* platelets (Fig 8D-F). Similar to a previously published report [42], Y-27632 did enhance PS exposure in wildtype platelets stimulated with thrombin (Fig 8G). The lack of effect on thrombin-stimulated *Trpc6^E896K/E896K^* platelets may relate to the already high degree of PS exposure under these conditions, though we cannot rule out that it is related to the modest effect of the inhibitor on the peak Fura2 ratio seen in *Trpc6^E896K/E896K^* platelets. Finally, Y-27632 also retarded clot retraction of wild-type PRP (Fig 8H). These results suggest a correlation between enhanced PS exposure and delayed clot retraction, whether mediated by gain-of-function TRPC6 or the ROCK inhibitor.

**Fig 8.**
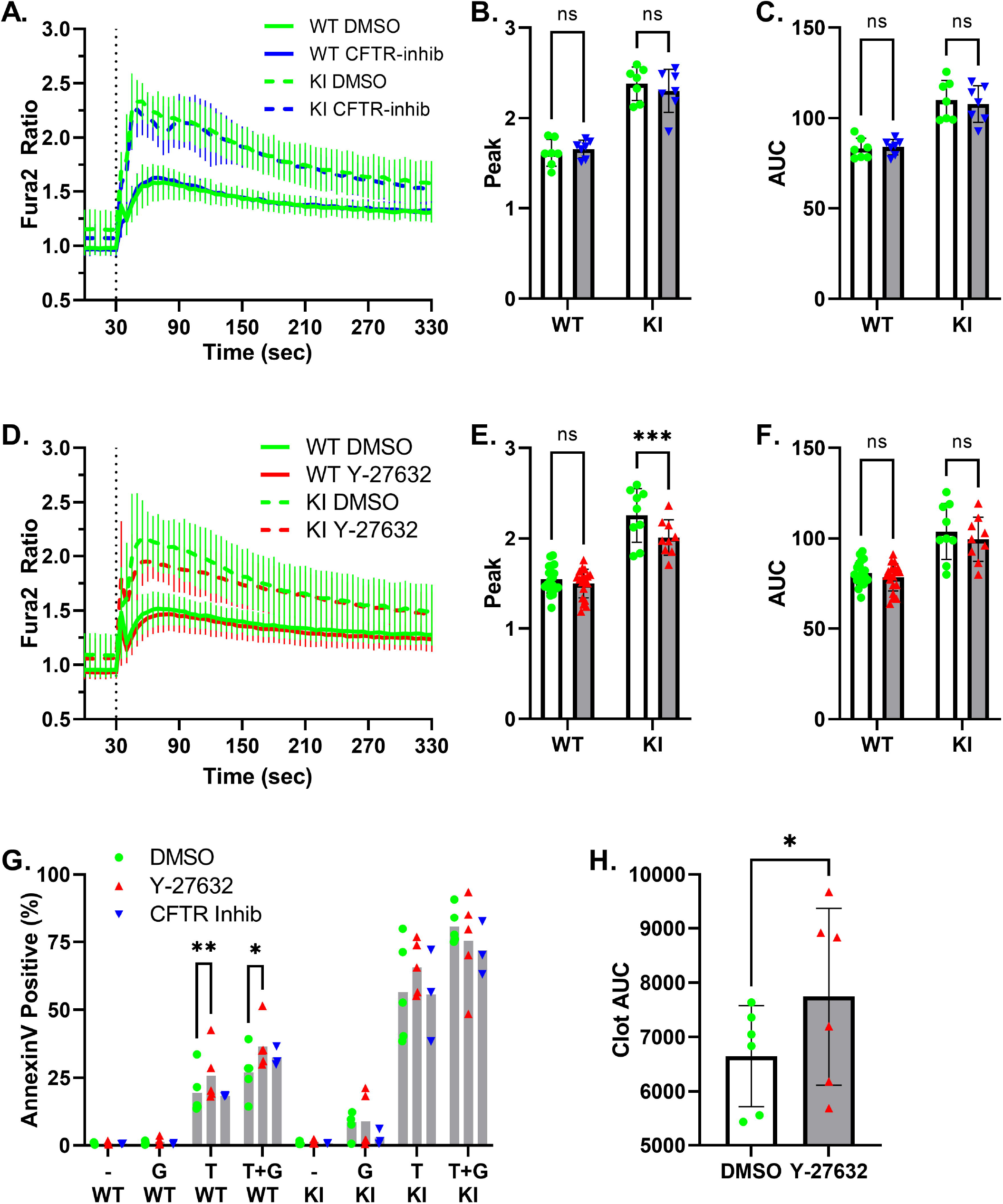
Effect of CFTR and ROCK inhibition on TRPC6-mediated platelet responses. A-C, WT and KI platelets, pre-incubated in the presence of vehicle (DMSO) or CFTR-inhibitor 172 (3 μM) for 15 minutes, were subject to Fura-2 ratiometric imaging. GSK was added after 30 seconds. A, Fura-2 fluorescence ratio (340/380) time-course. Shown are mean ± SD, n=7. The post-stimulation Fura-2 fluorescence ratio peak (B), and area under the curve (C), are shown as means and individual values. Two way ANOVA with Sidak’s multiple comparisons test. D-F, WT and KI platelets, pre-treated with ROCK inhibitor Y-27632 (10 μM) or vehicle for one hour, were subject to Fura-2 imaging and GSK stimulation. Shown are the time-course (D), peak ratio (E), and area under the curve (F); n=9-19. Two way ANOVA with Sidak’s multiple comparisons test. G, percentage of Annexin V stained platelets from WT and KI animals, pretreated with vehicle (DMSO), Y27632, or CFTR-inhibitor, followed by stimulation with vehicle (-), GSK (G), thrombin (T), or thrombin plus GSK (T+G); n=3-5. Mixed effects analysis with Dunnett’s multiple comparisons test; only statistically significant pairwise comparisons are shown. H, platelet-rich plasma was incubated with vehicle (DMSO) or Y-27632, and clot retraction observed. Clot area under the cure (AUC) was calculated by summing the clot areas across all time-points for each sample. Shown are means ± SD, and individual values; paired t-test.

## Discussion

By comparing the platelet phenotype of mice with wild-type, dominant negative, and gain-of-function *Trpc6* mutations, we provide further insight into the role of the TRPC6 channel in platelet function. Specifically, several conclusions can be drawn from this study. 1) TRPC6 is required for GSK1702934A-induced calcium influx in platelets, contributes a minor fraction of calcium influx downstream of thrombin stimulation, and is not significantly involved in calcium signaling downstream of ADP or thromboxane A_2_ receptor in these cells. 2) Wild-type TRPC6 is neither necessary nor sufficient for integrin αIIbβ3 activation, P-selectin surface exposure or PS exposure, but the TRPC6^E896K^ mutant can induce or enhance these processes. 3) In addition to calcium influx, some signaling responses show a graded response between *Trpc6* genotypes (e.g. Erk phosphorylation), whereas other signaling events (e.g. MLC2 phosphorylation, talin cleavage, clot retraction) are modulated exclusively downstream of mutant TRPC6^E896K^ channel. 4) Heterozygous *Trpc6^+/E896K^* platelets show a modest gain-of-function phenotype with regards to GSK-induced calcium influx, and thrombin-induced PS exposure, which does not extend to delayed clot retraction. Taken together, these results suggest that pathological processes mediated by gain-of-function TRPC6 may not be readily uncovered via extrapolation of differences between *Trpc6*-deficient and wild-type mice.

While the expression of TRPC6 in platelets is well established,[17–20] and confirmed here, there are conflicting accounts as to whether TRPC3 is also present and active in platelets.[18, 20] While we did not specifically investigate TRPC3 expression, we did find that GSK, reported to activate both TRPC3 and TRPC6,[43, 44] induced no appreciable Fura2 response in *Trpc6^-/-^* platelets. In FVB mice, the lack of a GSK response in *Trpc6^DN/DN^* and *Trpc6^+/DN^* platelets also argues that if TRPC3 is expressed, it does not form GSK-responsive channels independent of TRPC6.

We found that neither the presence of gain-of-function nor dominant negative TRPC6 impacted platelet integrin αIIbβ3 activation or degranulation in response to ADP, U46619 or thrombin. These results are consistent with prior studies reporting that loss of *Trpc6* does not influence these aspects of platelet activation.[18, 21] However, stimulating TRPC6 using GSK does augment the modest platelet activation induced by ADP. This suggests that as yet unidentified physiological activators of TRPC6 could enhance platelet activation synergistically with other weak platelet agonsists.

Influx of extracellular calcium is a well-established means of inducing phosphatidylserine exposure in platelets.[22] TRPC6, in conjunction with TRPC3, has previously been reported to be involved in PS exposure on platelets in response to simultaneous stimulation by thrombin and collagen.[20] In contrast, loss of both channels did not affect the small percentage of platelets exposing PS in response to thrombin, or collagen, stimulation alone. As we were unable to enhance PS exposure through the addition of either collagen or convulxin to thrombin (data not shown), we were unable to confirm the finding of Harper et al.[20] However, we found that dominant negative TRPC6 does not significantly impair thrombin-induced PS exposure, even though gain-of-function TRPC6 mutant enhances this process. In addition, TRPC6^E896K^ mutant is capable of inducing cleavage of talin, and delaying *in vitro* clot retraction, consistent with prior work demonstrating these pathways are activated by elevated intracellular calcium concentrations.[35, 36] We postulate that a certain threshold calcium influx is required to induce PS exposure and talin cleavage, and to impair clot retraction, which is attained by activating gain-of-function, but not wild-type, TRPC6. This would be consistent with recent structurefunction studies demonstrating that FSGS-associated mutations, including the E897K mutant, disrupt an inhibitory calcium-binding site in TRPC6.[45] We hoped to test this hypothesis with the use of TRPC6-PAM-C20, a positive allosteric modulator of TRPC6.[37] However, C20 has TRPC6-independent effects in platelets (as evidenced by inhibition of Annexin V staining of thrombin-stimulated *Trpc6^DN/DN^* platelets), precluding use of this strategy. We therefore cannot conclude whether the ability of TRPC6^E896K^ to induce PS exposure is solely due to the resultant increase in calcium influx, or whether additional mechanisms are involved.

In humans, gain-of-function TRPC6 mutations cause an incompletely penetrant, predominantly adult onset form of autosomal dominant FSGS.[14, 15] A platelet phenotype has not been reported. In contrast, Stormoken syndrome and York platelet syndrome, caused by dominant gain-of-function mutations in STIM1 or ORAI, and characterized by constitutive activation of store-operated calcium entry (SOCE), do. Hematologically, these disorders are characterized by a bleeding diathesis with thrombocytopenia, with platelets demonstrating evidence of activation, including PS exposure, in the absence of external stimulation.[46, 47] The lack of an overt platelet phenotype in humans with *TRPC6* mutations, and the subtle platelet phenotype in *Trpc6^E896K/E896K^* mice, suggests that TRPC6 plays a minor role in modulating calcium influx compared to SOCE. The fact that *TRPC6* mutations cause enhanced stimulated, but not basal, channel activity,[14, 28] unlike the STIM1 and ORAI mutations in Stormoken syndrome, could also contribute to these differences. Our results do suggest that platelet studies may be informative in characterizing the effects of human *TRPC6* mutations on endogenous channel activity and regulation. Whether abnormal platelet function contributes to the development of TRPC6-associated FSGS remains uncertain, though the lack of reported cases of disease recurrence after kidney transplantation argues against this.

In conclusion, while TRPC6 appears to be dispensable for most platelet functions, GOF mutations in this channel subtly alter platelet activation. Specifically, TRPC6^E896K^ mutant, but not wild-type TRPC6, channel can induce integrin αIIbβ3 activation, surface expression of P-selectin, phosphatidylserine exposure, myosin light chain phosphorylation and talin cleavage, and delay *in vitro* clot retraction. These results make clear that the pathological effects of GOF mutations may not be readily discernable by attempting to extrapolate from differences between wild-type and TRPC6-deficient model systems, but may represent true gain-of-function phenotypes. Future studies comparing *in vivo* platelet and hemostatic responses in the various murine *Trpc6* genotypes described here may help further elucidate the stimuli and signaling pathways that utilize this channel.

## Acknowledgements

We thank the Flow Cytometry, Small Animal Imaging, and Transgenic Cores at the Beth Israel Deaconess Medical Center, and the Harvard ICCB-Longwood Screening Core, for technical assistance and expertise. We are grateful to Martin Pollak and lab members for thoughtful discussions and suggestions.

Research reported in this publication was supported by the National Institute of Diabetes And Digestive And Kidney Diseases of the National Institutes of Health under Award Numbers R01DK115438 (to J.S.S.) and T32DK007726 (to B.R.S), and by the Klarman Scholarship Award (to J.S.S).

## Author Contribution

B. J. Brown and K. L. Boekell designed and performed experiments, and analyzed data. B. E. Talbot generated key reagents. J. S. Schlondorff designed and performed experiments, analyzed data, obtained funding, and wrote the manuscript.

